# Whole-brain Mechanism of Neurofeedback Therapy: Predictive Modeling of Neurofeedback Outcomes on Repetitive Negative Thinking in Depression

**DOI:** 10.1101/2023.11.16.567419

**Authors:** Masaya Misaki, Aki Tsuchiyagaito, Salvador M. Guinjoan, Michael L. Rohan, Martin P. Paulus

## Abstract

Real-time fMRI neurofeedback (rtfMRI-NF) has emerged as a promising intervention for psychiatric disorders, yet its clinical efficacy remains underexplored due to limited controls and an incomplete mechanistic understanding. This study aimed to elucidate the whole-brain mechanisms underpinning the effects of rtfMRI-NF on repetitive negative thinking in depression. In a double-blind randomized controlled trial, forty-three depressed individuals underwent NF training targeting the functional connectivity (FC) between the posterior cingulate cortex and the right temporoparietal junction, linked to rumination severity. Participants were randomly assigned to active or sham groups, with the sham group receiving synthesized feedback mimicking real NF signal patterns. The active group demonstrated a significant reduction in brooding rumination scores (p<0.001, d=-1.52), whereas the sham group did not (p=0.503, d=- 0.23). While the target FC did not show discernible training effects or group differences, we found that the interaction between brain activities during regulation and the response to the feedback signal was the critical factor in explaining treatment outcomes. Connectome-based predictive modeling (CPM) analysis, incorporating this interaction, successfully predicted rumination changes across both groups. The FCs significantly contributing to the prediction were distributed across broad brain regions, notably the frontal control, salience network, and subcortical reward processing areas. These results underscore the importance of considering the interplay between brain regulation activities and brain response to the feedback signal in understanding the therapeutic mechanisms of rtfMRI-NF. The study not only affirms the potential of rtfMRI-NF as a therapeutic intervention for repetitive negative thinking in depression but also highlights the need for a more nuanced understanding of the whole-brain mechanisms contributing to its efficacy.

## Introduction

Real-time fMRI neurofeedback (rtfMRI-NF) - a technique that enables participants to self-regulate functional brain activation - is considered a potential intervention for psychiatric disorders. Applied across various psychiatric conditions, this method’s feasibility for self-regulation of brain activation and its promising impact on symptom management are supported by numerous meta-analyses ^1–9^. Nevertheless, the clinical efficacy of rtfMRI-NF remains to be confirmed, as the majority of studies are preliminary, characterized by small sample sizes, the absence of stringent control conditions, and a partial understanding of the mechanisms of action. Overcoming these research gaps to ascertain the specific efficacy of NF treatments is an area of significant interest.

One strength of rtfMRI-NF is its ability to provide deep insights into whole-brain functional processes during self-regulation, extending beyond the targeted brain signal ^10^. Crucially, whole-brain analysis of rtfMRI-NF data is instrumental in decoding the mechanisms of action that underpin NF training. Emerging research has revealed diverse brain activities implicated in NF-mediated self-regulation, indicating that NF training encompasses a whole-brain process involving prefrontal control regions, the salience network, and reward processing areas ^11–14^.

Given that NF training might entail a reinforcement learning process ^15, 16^, examining brain activation in response to feedback signals is a valuable method for shedding light on the learning mechanisms driving self-regulation through NF. Investigations into brain responses to feedback signals have highlighted the pivotal roles of both prefrontal control regions and reward-responsive areas in NF-mediated self-regulation training ^14, 17, 18^. Significantly, the regions engaged during NF training and in response to feedback signals coincide with those involved in general skill acquisition ^19^ and emotion regulation tasks ^20, 21^. Moreover, these areas continue to be active even when a sham NF signal is administered, with participants unknowingly trying to control the feedback signal ^20, 21^. Consequently, these activations appear to represent attempts to modulate the feedback, irrespective of the training outcome. Therefore, pinpointing brain activations or connectivity patterns that foretell the success of training is still challenging.

The current study explores the whole-brain mechanisms underlying NF treatment, utilizing data from a prior double-blind, randomized controlled trial (RCT) ^22^. This earlier study demonstrated a significant decrease in symptoms of repetitive negative thinking exclusively in the active group following connectivity NF training, but not in the sham control group. Specifically, our study concentrated on the reduction of symptoms as the primary outcome of NF training, rather than on the self-regulation of the targeted brain signal. This approach contrasts with many studies that aim to elucidate the mechanisms of NF training, which have focused on identifying the neural substrates underpinning the acquisition of self-regulation of the target brain signal. The investigation into neural substrates of self-regulation may be justified if we assume that regulating the targeted brain signal facilitates changes in cognitive function or the symptoms of a disorder, thus rendering successful regulation essential for an effective intervention. However, this assumption is susceptible to the counterargument that other brain activities concurrently altered by training could be the drivers of behavioral changes, as posited by Kvamme, Ros ^23^. Because rtfMRI-NF training relies on the endogenous effort to self-regulate the signal and because participants cannot perceive the specific brain activation the feedback signal represents, the specificity of the NF intervention cannot be guaranteed. Kvamme, Ros ^23^ argue that we should not automatically assume a causal relationship between self-regulation of the target brain signal and the resultant behavioral outcomes. Corroboratively, research examining brain activations responsible for PTSD symptom amelioration post left amygdala upregulation training via rtfMRI-NF ^24^ revealed that activity in the dorsomedial prefrontal cortex (dmPFC) and middle cingulate cortex—not the amygdala—mediated symptom reduction. This indicates that the therapeutic effects may be attributed to brain activations beyond the intended target area.

Furthermore, symptomatic or behavioral improvements can occur independently of detectable changes in the targeted brain signal during training. Although the feedback signal is designed to reflect successful self-regulation, the desired training effect may not always be apparent in the targeted region, as evidenced by several neurofeedback studies. For example, Sukhodolsky, Walsh ^25^ utilized rtfMRI-NF to target the supplementary motor area (SMA) in adolescents with Tourette syndrome and noted significant symptom improvement in the active neurofeedback group, absent in the sham control group, despite a lack of significant SMA activation changes. Similarly, our earlier research ^22^ showed that neurofeedback aimed at enhancing the functional connectivity between the posterior cingulate cortex (PCC) and the right temporoparietal junction (rTPJ) markedly decreased rumination in depressed participants of the active group, with no corresponding change in the sham group, and without a notable difference in the targeted connectivity. Crucially, both studies implemented a stringent RCT methodology to ensure the effects were attributable to the neurofeedback. These results collectively imply that a comprehensive, whole-brain perspective may be essential to fully grasp the therapeutic mechanisms of rtfMRI-NF.

Consequently, the current study investigates the whole-brain mechanism of NF treatment by using data from our previous study ^22^. Our approach employs machine learning predictive modeling focusing on two distinct functional connectivity patterns: one during the self-regulation task and the other in response to neurofeedback signals. We use connectome-based predictive modeling (CPM) ^26^ to establish the relationship between these patterns and treatment outcome. By aggregating univariate features, multivariate predictive modeling techniques such as CPM can enhance both the sensitivity and robustness of predictions ^27–29^. We also examined the interplay between connectivity patterns during the task and in response to neurofeedback as a potential predictor of treatment success. Given the absence of significant differences in target connectivity between the active and sham groups and the consistent control of feedback signal amplitude, the key difference between the conditions may lie in the fidelity of the feedback to the participants’ actual brain activity. Thus, the interaction between brain activation and feedback signals could be critical in explaining the variance in symptom reduction observed between the groups.

## Methods

### Participants

We analyzed the data of the previous NF study for a treatment of repetitive negative thinking in depressed participants ^22^. The original study was registered on ClinicalTrials.gov (NCT04941066). In the study, forty-three individuals with Major Depressive Disorder (MDD) between 18 and 65 years of age were enrolled. The participants met the fifth edition of the Diagnostic and Statistical Manual of Mental Disorders (DMS-5) criteria for unipolar MDD. Further details of the inclusion and exclusion criteria were shown in Tsuchiyagaito, Misaki ^22^, and the CONSORT diagram is in the Supplementary Information (SI). All participants provided written informed consent. The study protocol was reviewed and approved by the WCG IRB (https://www.wcgirb.com) (IRB Tracking Number 20210286).

### Neurofeedback session procedures

Detailed procedures are described in the SI. Here, we provide an overview of the procedure. The participants were divided into the active (N=22) and sham (N=21) groups randomly. The active group received the NF of functional connectivity between the PCC and the rTPJ regions. The sham group received a feedback signal artificially synthesized to mimic the temporal probabilistic structure of the real NF signal in the active group.

Beyond employing an RCT design, the study offered several advantages for controlling the non-specific effects of neurofeedback training. Strict double-blinding was achieved by having a separate researcher remotely set up the experimental application, who did not interact with either the participants or the experimenter who engaged with the participants during the study. The sham neurofeedback signal was carefully designed to mimic the genuine neurofeedback signal in terms of reinforcement frequency and temporal pattern. The sham signal was controlled to avoid any unintended correlation with the target brain signal monitored in real-time. A post-session questionnaire confirmed that participants were unaware of their group assignment ^22^. Moreover, comprehensive real-time fMRI processing for noise reduction including physiological noise ^30^ ensured that the neurofeedback signal was free of confounding variables unrelated to brain activation, a critical consideration given that connectivity neurofeedback is known to be susceptible to physiological noise ^31^. The study confirmed that these confounding variables were effectively eliminated through comprehensive real-time fMRI processing ^32, 33^.

There were three consecutive NF training runs in the single session for each subject. Each NF training run was 8 m long with 90 s initial resting block, followed by 100 s regulation block with four consecutive presentations of negative trait words (25 s each) and a 30 s rest. The participants were engaged in the emotion regulation task (i.e., regulating negative thoughts while viewing the negative self-referential words) while receiving the connectivity NF. A positive feedback signal was presented when the target PCC-rTPJ FC reduced. The regulation and rest blocks were repeated three time in a run. Each participant also performed a baseline run and a transfer run where no feedback signal was presented; we only used the NF training runs for the prediction analysis.

### Connectome-based Predictive Modeling (CPM)

The detailed offline fMRI image preprocessing is described in the SI. After preprocessing, we calculated whole-brain functional connectivity using beta-series correlation ^34^. Specifically, we evaluated the beta values for the regressor of the regulation task for each NF regulation block using General Linear Model (GLM) analysis. We formed a series of beta values from the regulation task and used this series across blocks and runs to calculate the z-transformed Pearson correlation between each region in the Shen 268 atlas ^35^. Similarly, we estimated the betavalues for the response to the feedback signal for each block using GLM analysis (see SI) and calculated the beta-series correlation (z-transformed) between the same atlas regions. We then calculated the interaction between FCs (z-transformed correlations) for the regulation task (RegTask) and the response to the NF signal (RespNF) by multiplying their normalized values. It is important to note that the NF signal reflects approximately 6 to 8 seconds of the previous brain state due to hemodynamic response delay and fMRI signal acquisition time. As a result, TR-wise interaction between RegTask and RespNF connectivity cannot indicate the contingency between these responses. Our current approach, which decomposes the signal into block-wise series, helps mitigate this issue at the expense of temporal resolution.

We employed Connectome-Based Predictive Modeling (CPM) analysis to identify brain activation patterns during neurofeedback (NF) training that could predict treatment outcomes. The target value for prediction was the change ratio of the Ruminative Response Style Brooding Subscale (RRS-B) score ^36^, measured one week after the NF training session, relative to the baseline. We focused on the brooding subscale because it specifically leads to worse prognosis ^37^. We used the whole-brain FC patterns of the regulation task (RegTask), the response to the NF signal (RespNF), and their interaction (RegTask:RespNF) individually to construct the CPM model using various combinations. Specifically, we evaluated five different CPM models: 1) RegTask, 2) RespNF, 3) RegTask + RespNF, 4) RegTask + RespNF + RegTask:RespNF, and 5) RegTask + RegTask:RespNF. We included model 5 because we found that the RespNF term alone did not provide significant predictive information. The model with RegTask:RespNF alone was also tested as a post-hoc evaluation.

Predictive performance was assessed using 5-fold cross-validation with covariate (age, sex, head motion) regression and hyperparameter optimization in the nested cross-validation. The process was repeated 100 times with different random splitting of the validation set to obtain a confidence interval and a reliable estimate of predictive performance. Further details of the model training procedures are described in the SI.

### Statistical analyses

The statistical significance of CPM performance was evaluated using a permutation test. The output values (ratio of change in RRS-B relative to baseline: dRRS-B) were randomly permuted 1000 times, and in each permutation iteration, 5-fold cross-validation was repeated 10 times with different random splits. The same covariate regression and hyperparameter optimization procedure including the nested cross-validation was also applied during the permutation test. Note that the covariate regression for the dRRS-B was performed for the non-permutated data, since the test needs to evaluate the null distribution of the model performance apart from the covariate effects (see Winkler, Ridgway ^38^ for further discussion). The median of 10 replicates was taken in each iteration to generate a null distribution.

Symptom change in RRS-B was tested using linear mixed effect (LME) model analysis with the fixed effects of session (pre/post), group (active/sham), age, and sex, and the random effect of participant to the intercept. The lme4 package ^39^ with the lmerTest package ^40^ was used, and each contrast was calculated using the emmeans package ^41^ in the R language and statistical computing ^42^. All the *p* values reported in the post-hoc contrast analysis were corrected using multivariate *t* distribution^41^. In these analyses, *p* < 0.05 (corrected as necessary) was considered significant.

## Results

### Data selection

The participants with more than 30% censored TRs due to head motion (frame-wise displacement [FD] > 0.3 mm) were excluded from the analysis. We further excluded the blocks with more than 30% censored TRs within the task block and the participants with less than five remaining blocks were also excluded from the analysis. There was no significant difference in the number of dropped blocks between groups (*t*(34)=1.500, *p*=0.143). As a result, the number of analyzed participants were 16 (11 females, mean age 33.5) for the active group and 18 (13 females, mean age 33.5) for the sham group. For the selected participants, there were no significant differences in age (*t*(32)=0.002, *p*=0.987) and sex composition (*χ*^2^(1)=0.049, p=0.825) between the groups.

### Significant treatment effect on RRS-B for the active group alone without changes in the NF target FC

Significant reduction in RRS-B score was observed for the active group but not for the sham group. Figure 1 shows the mean and each participant’s RRS-B scores at the pre and post-training sessions for each group. While the main effect of group was not significant (*F*[1,30]=0.062, *p*=0.805), the main effect of session (*F*[1,32]=12.944, *p*=0.001) and the interaction between the session and group (*F*[1,32]=7.125, *p*=0.012) were significant with LME analysis. The post-hoc analysis indicated that the post-pre difference was significant for the active group (*t*[32]=-4.31, *p*<0.001, *d*=-1.52) but not for the sham group (*t*[32]=-0.67, p=0.503, *d*=-0.23).

**Figure 1.**
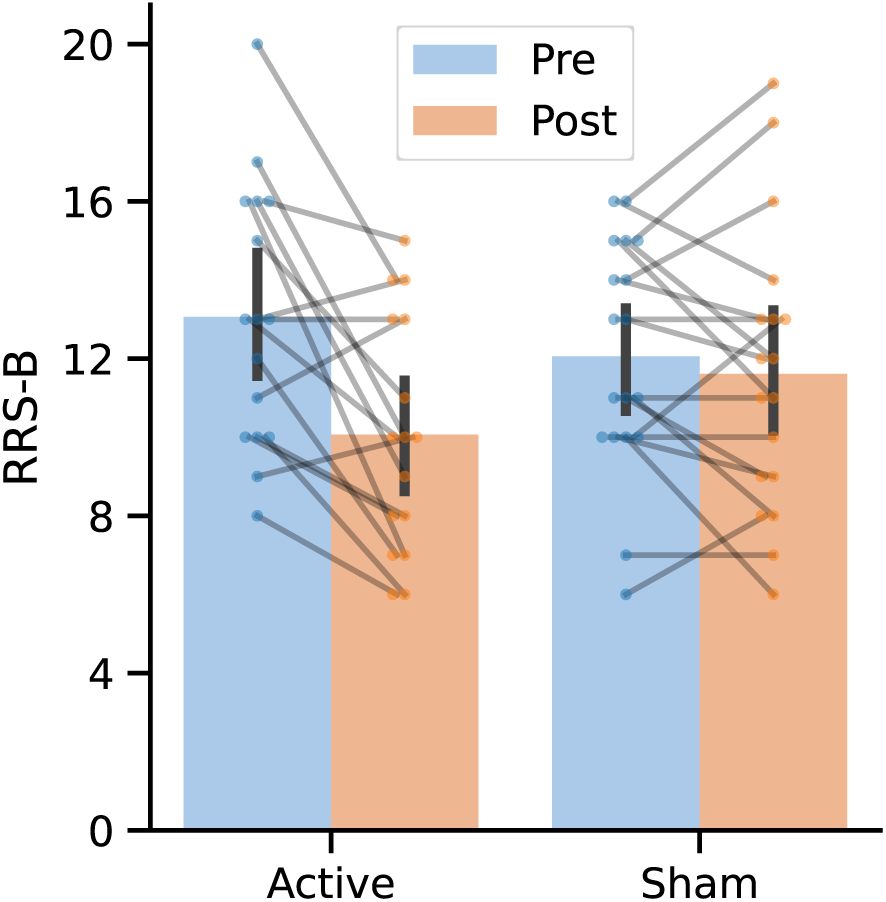
Rumination scores before (pre) and after (post) NF training for both the active and sham groups. The bar plots represent the mean, and the error bars depict the 95% confidence interval. Individual participants are denoted by the points, with connecting lines between them. RRS-B: Ruminative Response Style, Brooding subscale.

In contrast to the significant treatment effect on the target symptom score, there was no significant effect on the NF target, PCC-rTPJ FC. Figure 2 shows the psychophysiological interaction (PPI) beta values for the PCC seed region (x, y, z = −6, −58, 48 mm) to the rTPJ (51, - 49, 23 mm) region across training sessions (seed and the target regions were defined by 6mm-radius sphere). An LME analysis with the fixed effects of run, group, age, sex, head motion (mean FD), and the random effect of participant for the intercept showed no significant main effect of the run (*F*[4, 478]=0.917, *p*=0.454), group (*F*[1, 37]=0.104, *p*=0.748), and run by group interaction (*F*[4, 478]=0.633, *p*=0.639) on the NF target connectivity.

**Figure 2:**
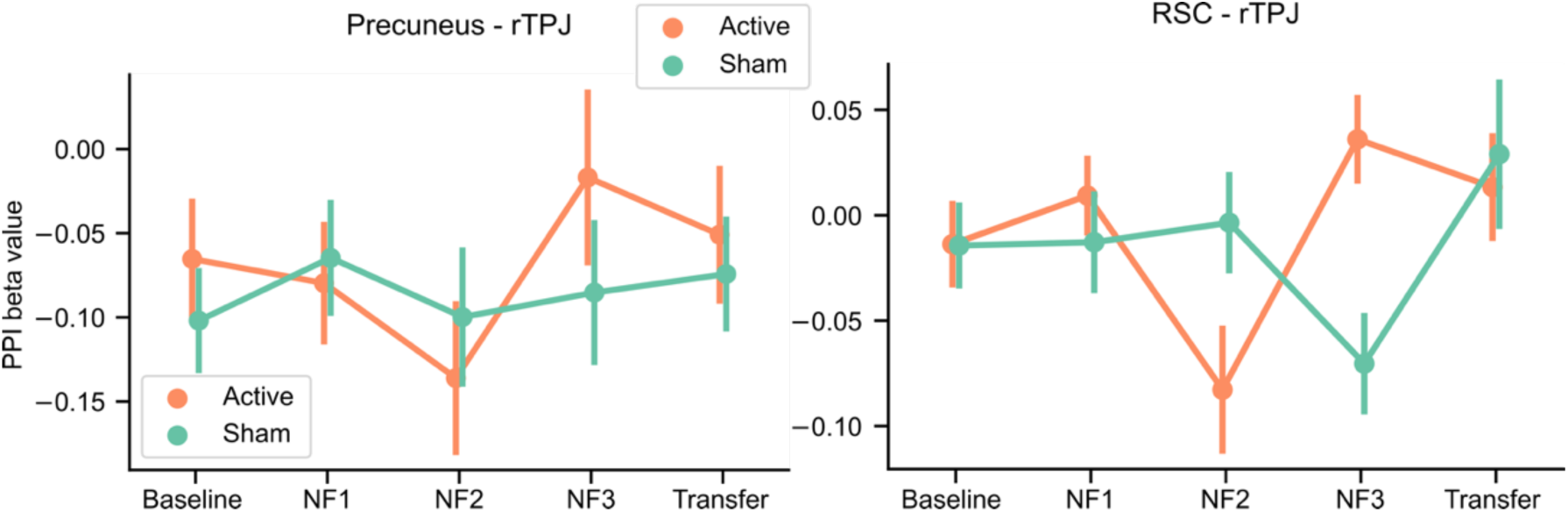
Results of the PPI analysis (beta value) for the target (PCC-rTPJ) and proximal RSC (RSC-rTPJ) functional connectivity across runs. PCC: posterior cingulate cortex, rTPJ: right temporoparietal junction, RSC: retrosplenial cortex.

Additionally, Tsuchiyagaito, Misaki ^22^ identified the other brain region in the retrosplenial cortex (RSC, [−7, −53, 11 mm]) that showed a different training effect between the active and the sham group. For the presently selected data, the PPI analysis between the RSC-rTPJ revealed no significant main effects of the run (*F*[4, 478]=2.178, *p*=0.070) and group (*F*[1, 29]=0.216, *p*=0.645), but significant run by group interaction (*F*[4, 477]=3.885, *p*=0.004). Post-hoc analysis indicated that the group difference was significant at NF2 (active < sham, *t*[334] = 2.354, *p*=0.019, *d*=0.475) and NF3 (acitve > sham, *t*[352]=-2.958, *p*=0.003, *d*=-0.637). However, when we calculated the Spearman correlation between the mean RSC-rTPJ beta values in the NF1, NF2, NF3 runs and the RRS-B change, no significant correlation was observed (*rho*=-0.035, *p*=0.843), indicating that the RSC-rTPJ connectivity during the NF training was not predictive of symptom change.

### CPM analysis

Figure 3 shows distributions of CPM analysis results for each model specifications. The median prediction performance was not significant for the CPM models with RegTask (median=0.194, *p*=0.988), RespNF (median=0.012, *p*=0.996), and RegTask+RespNF (median=0.135, *p*=0.990). In contrast, the models with the interaction term RegTask+RespNF+RegTask:RespNF (median=0.190, *p*=0.001) and RegTask+RegTask:RespNF (median=0.272, *p*<0.001) showed significant prediction performances. The performance difference between the models with and without RespNF main effect term was significant (*p*=0.007) with better result by excluding the RespNF term. We also evaluated the CPM only with the interaction term, but its performance was significantly worse (*p*=0.007) than the model with the main effect of RegTask. Thus, the model with the FC patterns during the regulation task (RegTask) and its interaction with the FC in response to the feedback signal (RespNF) was the best to predict the RRS-B symptom change.

**Figure 3.**
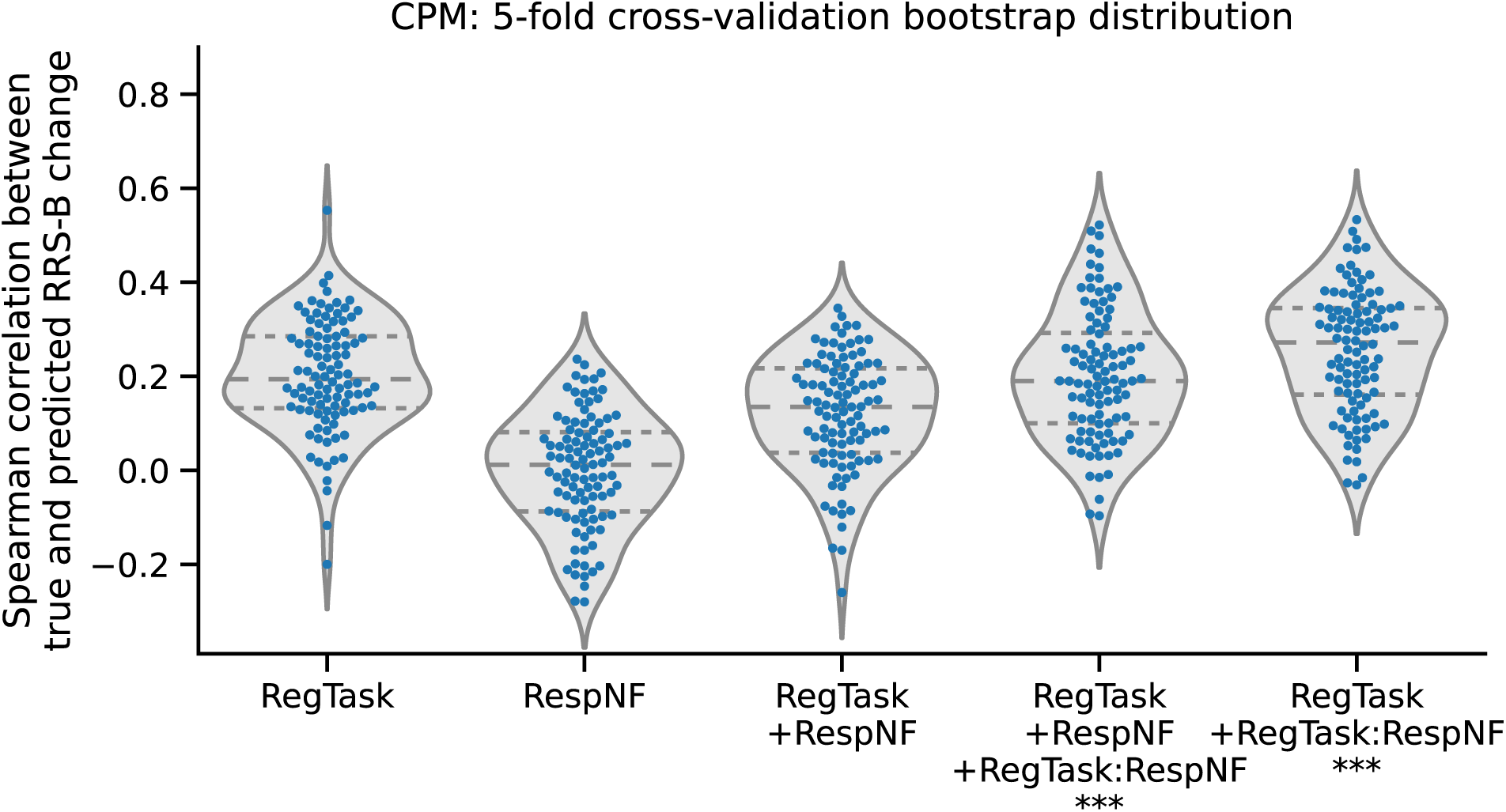
Distributions of the CPM prediction performance (as measured by the Spearman correlation between true and predicted values) for the RRS-B score change following NF training, compared to the baseline assessment. Each point represents a single iteration from the 5-fold cross-validation results (with a total of 100 iterations using different random splits). The violin plot illustrates the distribution curve, and the horizontal lines indicate quartile positions. ***: *p* < 0.001.

We further examined the connectivity that contributed to the prediction for the best performing model. Figure 4 shows the connectivity that were selected 76% times across 100 bootstrap model trainings (*p* < 5e^-8^ using a binominal test), which corresponds to *p*<0.001 with Bonferroni correction across all the 27028 connectivities. The plots indicate that many connectivities remained in spite of this stringent threshold, indicating that the model consistently employed broadly distributed connectivity for the prediction. While connectivities that informed prediction distributed across the brain, several dense connectivity regions were also observed in the circle plot. Supplementary Tables S1 and S2 show the top 10 nodes with the highest sum of absolute correlation with RRS-B reduction across their significant connectivities. For the RegTask, predictive connectivity was dense in the cingulo-opercular task control regions, salience network regions, and the subcortical thalamus and basal ganglia regions. Similarly, for the RegTask:RespNF interaction, predictive connectivity was dense in the fronto-parietal task Control regions, salience network regions, and the subcortical hippocampus, thalamus, basal ganglia regions.

**Figure 4.**
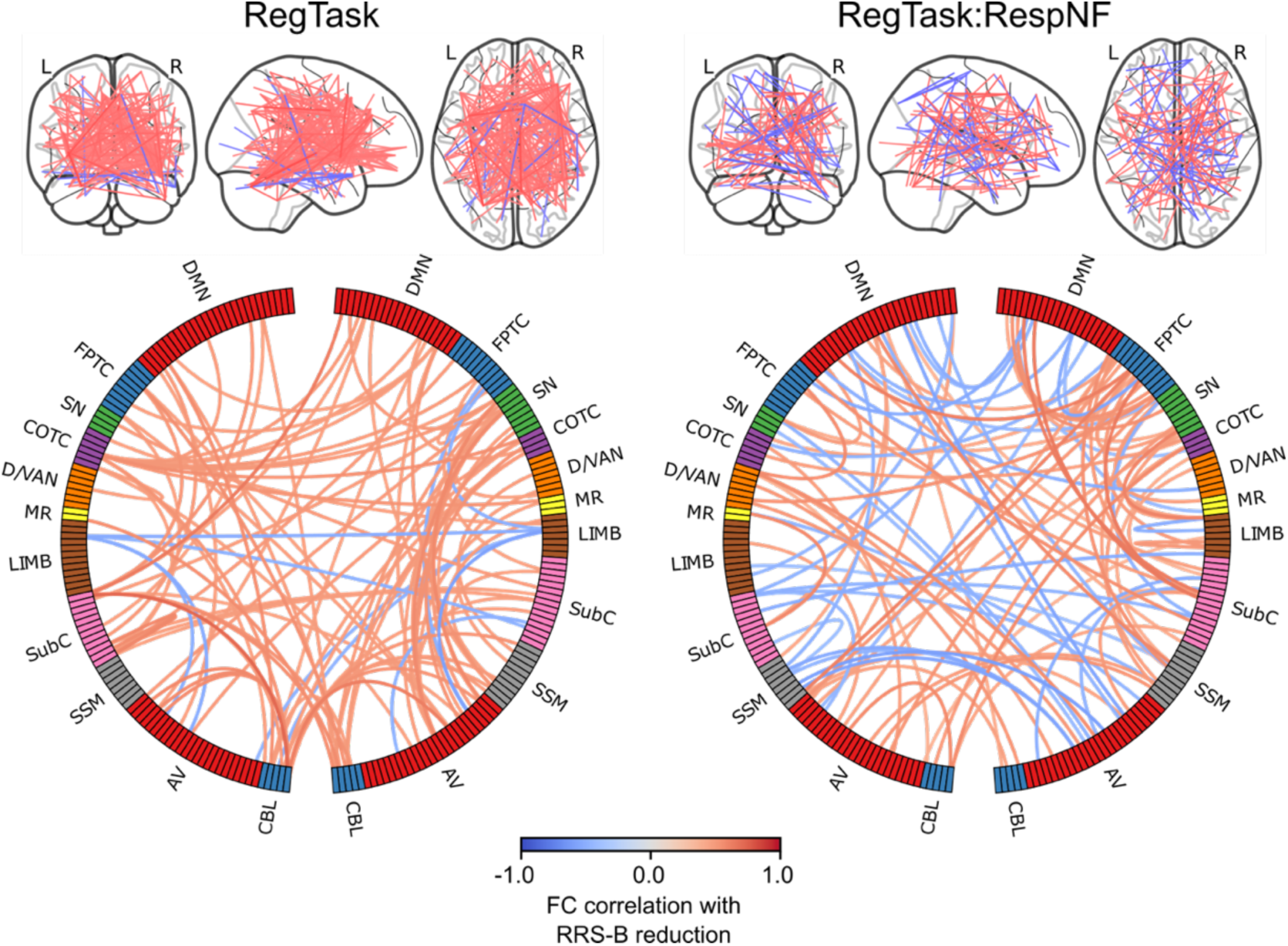
Plots of the connectivity selected by the CPM model in more than 76% (*p* < 0.001 with Bonferroni correction) of the cross-validation iterations. Line color indicates the connectivity correlation with RRS-B. Warm color indicate that the higher the FC the more RRS-B decreased, and the cool color was vice versa. The circle plots summarized for each network region. Network labels are adapted from Drysdale, Grosenick ^47^. DMN: Default Mode Network, FPTC: Fronto-Parietal Task Control, SN: Salience Network, COTC: Cingulo-Opperculum Task Control, D/VAN: Doral Visual Attention Network, MR: Memory Retrieval, LIMB: default mode/limbic, SubC: Subcortical, SSM: Sensory SomatoMotor, AV: Auditory-Visual, CBL: Cerebellum.

Figure 5 illustrates the relationship between the mean connectivity (z) for the informative FCs in the prediction model and the changes in RRS-B. These plots are intended to investigate the potential confounding effect of group differences on the predictions, rather than to present statistical analysis of these relationships, which could constitute double-dipping since the FCs correlating with RRS-B changes were selected for the CPM. The plots show that group differences did not affect the prediction, confirming that the selected FCs effectively predict changes in RRS-B scores in both active and sham groups.

**Figure 5.**
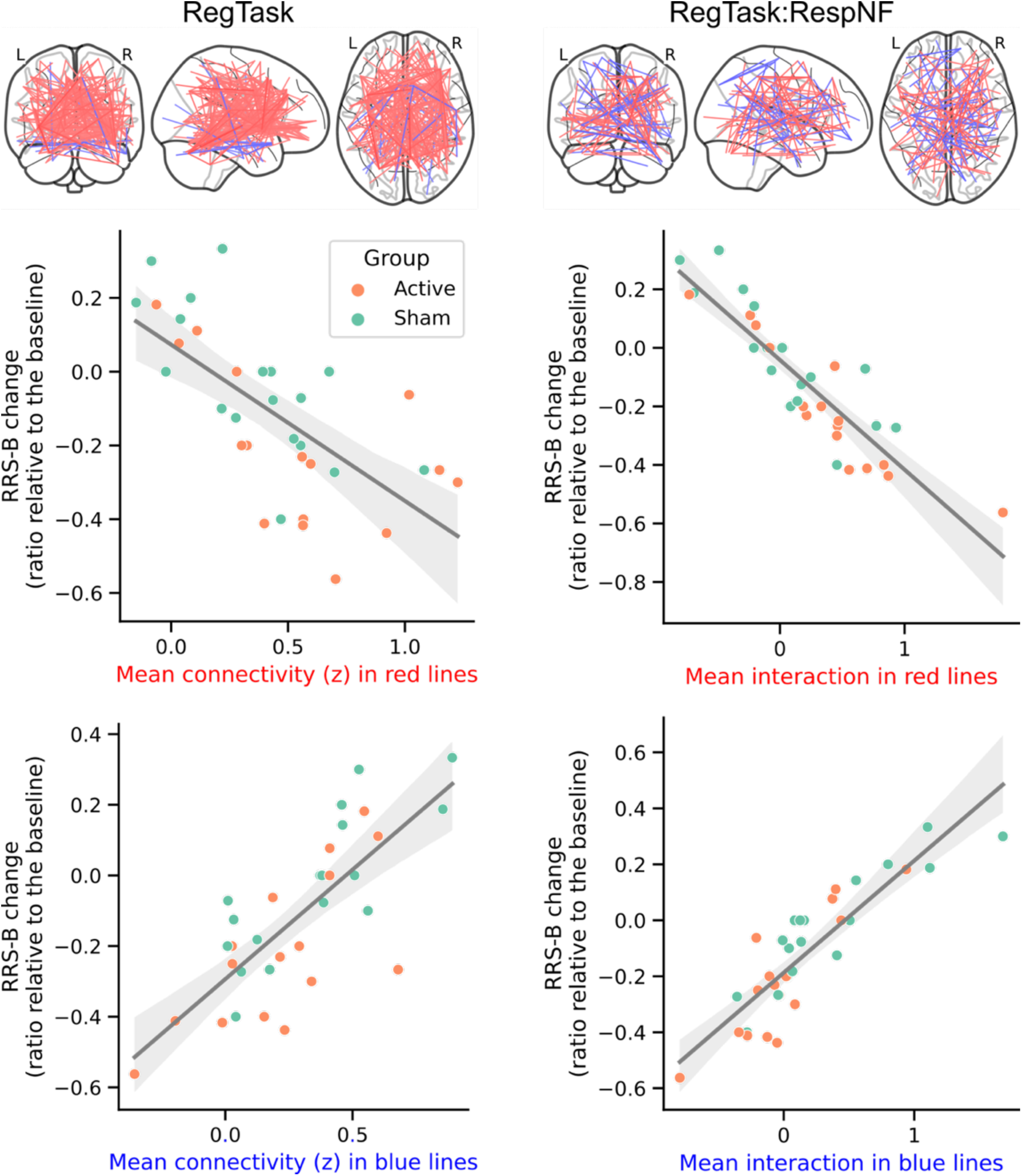
Association between the RRS-B change and the mean connectivity values in the informative FCs for CPM prediction, both for positively and negatively related FCs during the regulation task (RegTask) and its interaction with the response to the neurofeedback signal (RespNF). The line represents the fitted trend, and its shadow indicates the 95% confidence interval.xs

### Association of other factors with RRS-B changes

We also examined other possibly predictive factors of RRS-B change, including age, sex, head motion, self-rating of the regulation success, the duration of the positive feedback presentation, the interaction between the self-rating of the regulation success and the group, and the interaction between the positive feedback duration and the group. However, linear model analysis with these factors on the RRS-B change (ratio relative to the baseline) indicated no significant effects in any of these (refer to SI for details).

### Voxel-wise activation analysis for the association with RRS-B change

Additionally, we conducted a voxel-wise mass univariate analysis on both the RegTask and RespNF beta maps, as well as on their interaction map, to determine if voxel-wise activation linked with treatment effects was reflected in region-wise responses. Detailed results of this analysis are provided in the SI. We observed a significant association between symptom change and RegTask activation only. The regions involved were different from the NF target areas, but were seen in the lateral occipital, superior frontal, precentral gyrus, and thalamus regions. These regions consistently show a decrease in their activation associated with RRS-B reduction, suggesting that the effects in these regions may be due to a reduction in task load over successive runs (detailed results are available in the SI).

## Discussion

The primary objective of this study was to investigate the whole-brain mechanisms underlying the effects of rtfMRI-NF on repetitive negative thinking, specifically brooding rumination (RRS-B), in depressed individuals. Our results demonstrated a significant reduction in RRS-B scores in the active neurofeedback group but not in the sham group, although no significant changes were observed in the targeted FC between the PCC and the rTPJ. Interestingly, CPM analysis revealed that the most effective models for predicting RRS-B reduction incorporated both the FC patterns during the regulation task (RegTask) and their interaction with the FC patterns in response to the feedback signal (RespNF). These models employed broadly distributed connectivity across multiple brain regions, including the cingulo-opercular task control and salience network regions. Other potentially predictive factors such as self-rating of regulatory success and the duration of the positive feedback signal showed no significant effects on RRS- B changes. In summary, our findings suggest that the efficacy of rtfMRI-NF in reducing brooding rumination is not solely dependent on the modulation of targeted FC but involves a complex interplay of whole-brain connectivity patterns, thereby challenging the traditional focus on targeted brain regions in neurofeedback research.

In agreement with our hypothesis, connectivity associated with the reduction in rumination was not localized to a specific region but was widely distributed across the brain, primarily in the frontal control, salience network, and subcortical reward processing areas. This finding is consistent with a meta-analysis of rtfMRI-NF training studies across different target regions ^11^. The meta-analysis found involvement of a broad array of brain regions during NF training, including executive control regions (i.e., ventrolateral prefrontal cortex [vlPFC], dorsolateral prefrontal cortex [DLPFC], premotor cortex), salience network regions (i.e., anterior insula, anterior cingulate cortex [ACC]), reward processing area (i.e., striatum), temporo-parietal areas, lateral occipital regions, and the temporo-occipital junction bilaterally. Investigations into functional connectivity with target brain regions like the amygdala have also reported increased connectivity with prefrontal areas, including the DLPFC, dorsomedial and ventromedial PFC, as well as the ACC ^12–14^.

Studies examining responses to feedback signals during rtfMRI-NF training have also demonstrated the involvement of these regions. Lawrence, Su ^17^ examined brain activation correlated with the amplitude of the feedback signal during rtfMRI-NF training aimed at increasing right anterior insula activation. They observed a positive correlation with the feedback signal in the dorsal ACC and left supramarginal gyrus, and a negative correlation in the inferior and rostral regions of the ACC and primary visual cortex. Similarly, Paret, Zahringer ^14^ analyzed brain activation in response to the NF signal during up- and down-regulation training of the right amygdala. They found that the medial thalamus was active in monitoring feedback signals broadly, while the ventral striatum (VS) responded specifically to the reinforcement signal, with VS activation showing a positive correlation with amygdala activation. Skottnik, Sorger ^18^ also showed that self-regulatory performance was correlated with striatal activity. The involvement of these regions is consistent with previous research highlighting their role in emotion regulation ^20, 21^ and skill learning ^19^. Taken together, these findings suggest that both prefrontal control regions and reward-responsive areas play an important role in NF-mediated self-regulatory training.

However, a previous study also suggested that these regions were activated during sham NF and were not necessarily indicative of training success ^43^. Our results suggest that effective treatment occurs when participants not only activate these emotion regulation and skill learning networks, but also receive consistent feedback that can guide them toward a desired neural state. Given that the interaction between RegTask and RespNF was critical in predicting treatment effect, participants who activate these networks in response to the feedback signal in a manner consistent with regulatory activation are more likely to experience a successful treatment outcome. Thus, effective treatment outcomes may depend on how individuals adapt and regulate their brain activation in response to neurofeedback, rather than simply trying to regulate it.

The whole-brain distributed associations also imply that the efficacy of NF treatment does not depend solely on the regulation of the target brain signal. Instead, this target signal may act as a guide, directing the brain toward a desired state. If this hypothesis proves valid, we may need to rethink our strategy for selecting the NF target signal. Traditionally, the NF target region has been identified based on its relationship to the target cognitive function or symptomatic state, assuming a causal relationship between the brain signal and the symptomatic state. However, the primary criterion for selecting a target region for the NF signal may not be its direct causal relationship with the symptoms. Instead, the key factor should be its ability to accurately represent the related cognitive and symptomatic states. In our current study, the PCC-rTPJ FC was used as the NF target. This FC remained low throughout the NF sessions (see Figure 2), with no significant variation observed across sessions. Therefore, on average, participants successfully regulated the NF signal in the instructed direction. As this NF signal has been associated with rumination in depression ^33^, it could have served as an indicator of the brain state associated with reduced rumination.

Notably, the present CPM analysis predicted RRS-B reduction in both the active and sham groups. Some participants in the sham group who experienced a decrease in RRS-B had FC values similar to those of active group responders, as shown in Figure 5. These findings suggest that responders in the sham group may be undergoing an adaptive training process similar to that of the active group. Pecina, Chen ^44^ noted that individuals could selectively respond to positive feedback while disregarding negative feedback, influenced by treatment expectations. Hence, even random NF signals might be leveraged to adjust brain activity towards a desired state. Nevertheless, our results, showing a significant difference between active and sham groups, underline the necessity of an accurate NF signal for effective brain state modulation, while also highlighting the complexity of placebo effects in NF training ^44–46^. Several limitations of the current study merit discussion. Firstly, our results highlight the interaction between FC patterns during the regulation and the response to the NF signal as predictive of subsequent treatment effects on rumination. However, the NF signal reflects brain activations from several seconds prior, due to the inherent delay in the hemodynamic response and the time required for fMRI imaging. This makes establishing a direct link between regulation activity and the NF signal response challenging. To address this, we analyzed prolonged block-wise responses and calculated beta-series correlations to determine FC, but this method sacrifices temporal resolution and inhibits the assessment of learning effects over multiple runs due to limited data blocks. Future research should design experiments that tackle these challenges while enabling temporal analysis. Another significant limitation is the small sample size, necessitating caution in drawing definitive conclusions. While our findings are promising, they require validation through larger cohort studies. We employed strict motion thresholds to mitigate potential confounding effects on symptom reduction. However, this stringent criterion led to a reduced sample size and an attrition rate of approximately 20%. Future studies should anticipate and account for such attrition rates.

In conclusion, this study sought to explore the whole-brain mechanisms underlying the effects of rtfMRI-NF on brooding rumination in depression. The results showed that treatment efficacy can be predicted by broad patterns of whole-brain FC. In particular, the interaction between regulatory activity and response to the neurofeedback signal emerged as a pivotal predictor of treatment outcome. This suggests that while rtfMRI-NF may focus on a specific area, its effects are distributed, impacting an extensive network of brain regions. Our findings highlight a potential oversight in studies that assume direct changes in targeted signals are necessary for behavioral improvement, potentially overlooking broader effects of NF. As the field of NF research advances, it will be critical to broaden outcome measures to include changes in cognitive function or symptomatology. Furthermore, comprehensive whole-brain analysis is essential to fully understand the complex neurophysiological changes induced by NF training.

