## Supplementary material for "Whole-brain Mechanism of Neurofeedback Therapy: Predictive Modeling of Neurofeedback Outcomes on Repetitive Negative Thinking in Depression": SI

### CONSORT diagram for participants enrollment and selection

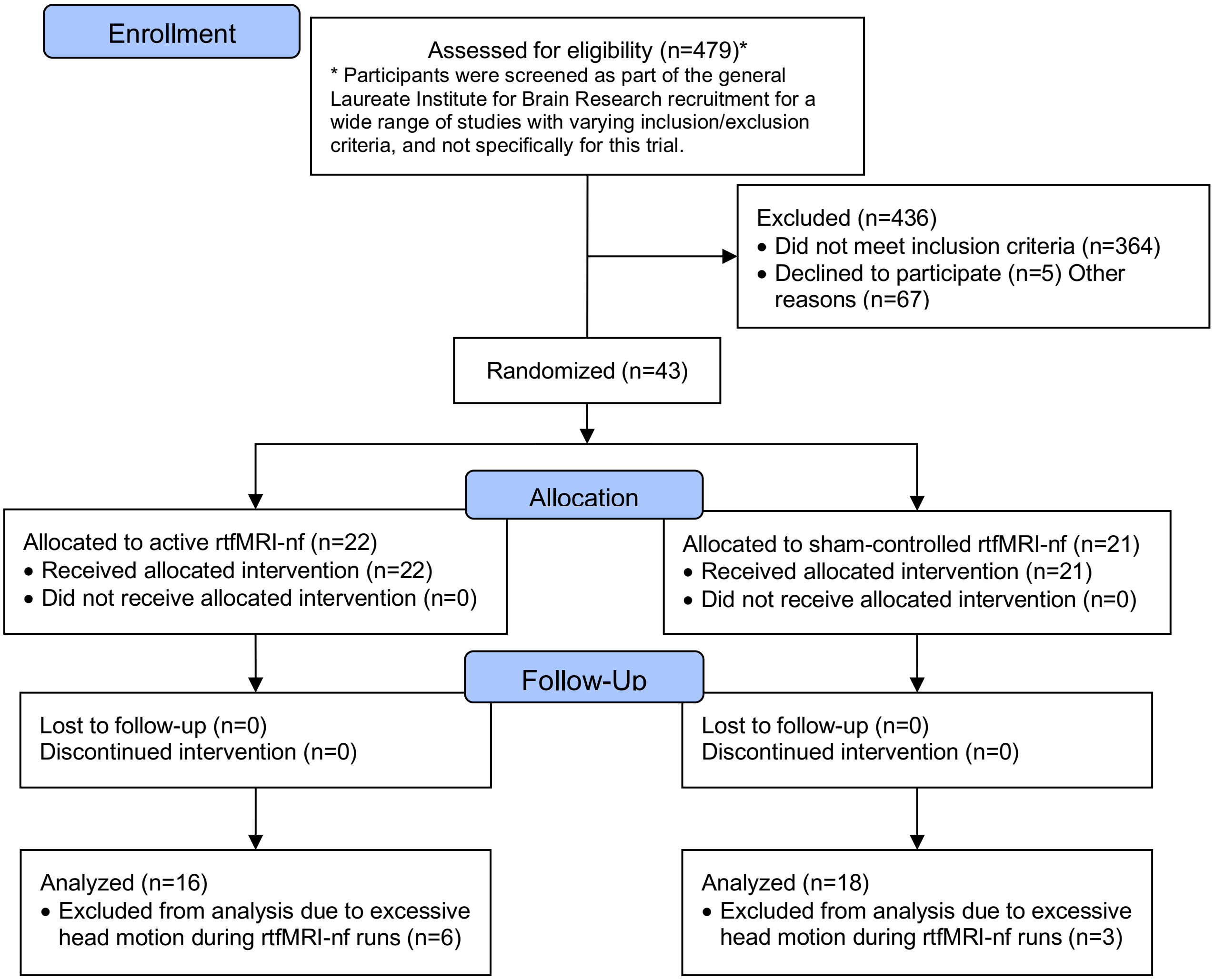

### Neurofeedback session procedures

Participants were randomly assigned to the active (N=22) and sham (N=21) groups. We confirmed that participants' reports of guessing their group assignment were independent of the actual assignment [1]. The active group received neurofeedback (NF) of functional connectivity between the posterior cingulate cortex (PCC) and the right temporoparietal junction (rTPJ) regions, which was correlated with the brooding subscale of the Ruminative Response Style [2] (RRS) score [3]. We focused on the brooding subscale (RRS-B) because it specifically leads to a worse prognosis [4]. The functional connectivity NF signal was calculated with the two-point method [5] and presented at each TR during the regulation task block. In the two-point method, the positive feedback signal was presented when the signals of the two regions (i.e., PCC and rTPJ) changed in different directions to train participants to reduce the target FC.

The sham group received a feedback signal that was artificially synthesized to mimic the probabilistic structure of the real NF signal in the active group. The sham feedback signal had the same gain frequency and temporal transition pattern as the real NF signal. The probabilistic structure was updated each time a new active participant was enrolled, and the initial parameters were evaluated from the results of the healthy participants in a previous study [6]. Thus, the order of enrollment of the active and sham groups was blinded to the experimenter handling the participants. In addition, we calculated the correlation between the time courses of the synthesized sham signals and the actual target connectivity measured in real time during the scan, and when the absolute correlation exceeded the threshold (> 0.3), the feedback signal was made to reduce the absolute correlation. This procedure ensured that the sham signal was safely independent of the target brain activity.

One NF training run was 8 m long with an initial rest block of 90 s, followed by a 100 s regulation block with four consecutive presentations of negative feature words (25 s each) and a 30 s rest. The regulation and rest blocks were repeated three times in one run. Each participant completed three neurofeedback training runs between the baseline and transfer runs, in which no feedback signal was presented. Participants were engaged in the emotion regulation task (i.e., regulating negative thoughts while viewing the negative self-referential adjective word) while receiving connectivity NF. The feedback signal is presented by blue bars displayed to the side of the adjective word when the signals of the two regions (i.e., PCC and rTPJ) changed in different directions to train participants to reduce the target FC [5, 6].

Real-time fMRI NF was performed on a 3T MR750 Discovery MRI (GE Healthcare, Milwaukee, WI). Functional BOLD contrast images were acquired using a T2*-weighted gradient echo planar imaging with TE/TR=25ms/2s matrix 96x96 on a 240 mm FOV, 40 slices at 2.9 mm, SENSE = 2. The fMRI signal for NF was subjected to a comprehensive real-time noise reduction process using the RTPSpy library [7]. This included slice timing correction, motion correction, spatial smoothing with a 6 mm FWHM Gaussian kernel within the brain mask, scaling to a percentage change relative to the average of the first 28 TRs (in the initial rest period), and regressing out noise components. The noise regressors were 12 motion parameters (three shifts, three rotations, and their temporal derivatives), eight RETROICOR regressors (four cardiac and four respiratory), global signal, white matter mean signal, ventricular mean signal, and Legendre polynomial models of slow signal fluctuations. While the connectivity NF signal is susceptible to physiological and motion noise in rtfMRI-NF [8], we confirmed that our real-time connectivity NF signal was free of physiological and motion noise after this processing [7, 9]. These careful controls of the NF signal ensured that the NF signal in the active group reflected their real brain activation with minimal noise effect, and the sham group received the same treatment with the same NF gain probability as the active group, except that the NF signal was not correlated with their brain activation.

### Offline fMRI image processing and block-wise GLM analysis

We used Analysis of Functional NeuroImage (AFNI; http://afni.nimh.nih.gov/) for offline fMRI image preprocessing. The process included discarding the first three TRs to await steady-state, despike, RETROICOR, and respiratory volume per time (RVT) corrections, slice timing and motion corrections, nonlinear warping to the MNI template brain with resampling to 2 $mm^3$ voxels using the ANTs (http://stnava.github.io/ANTs/), spatial smoothing with a 6 mm FWHM Gaussian kernel, and scaling the signal to the percent change relative to the mean in each voxel.

General linear model (GLM) analysis was used to evaluate a response (beta coefficient) for each block of NF regulation. We used a least squares - separate (LS-S) approach [10] in which a regressor for a target block and a regressor for all other blocks were entered into the design matrix to obtain the estimate for one block. This was repeated for each block in an independent GLM analysis. This approach can avoid collinearity problems when block-wise regressors for all trials are entered into a GLM design matrix. It has been suggested that this approach can provide a better estimate of the block-wise response [10]. The GLM was run using the AFNI 3dREMLfit command. Each time point with large motion (> 0.3 mm frame-wise displacement [FD]) was censored within the GLM, and the design matrix included noise regressors of 12 motion parameters (3 shifts, 3 rotations, and their temporal derivatives), three principal components of the ventricular signals, event-related response model for the word change, and the local white matter average signal (ANATICOR [11]).

The response to the feedback signal was also estimated with the GLM analysis. The neurofeedback was the binary signal that was turned on when the PCC and rTPJ signals moved in opposite directions and turned off when the signals moved in the same direction. The regressor of the NF signal response was a boxcar function indicating the presentation of the positive feedback signal, convolved with a canonical hemodynamic response function, and orthogonalized to the task block regressors to avoid the confounding effect of the task block response. The NF response was evaluated for each block using the LS-S approach [10].

### Whole-brain functional connectivity calculation

Using the series of beta estimates of each NF regulatory block across runs, whole-brain functional connectivity was calculated using the correlation of the beta series [12]. Specifically, the mean beta values in the Shen 268 atlas regions [13] were calculated and the z-transformed Pearson correlation between the beta series of the atlas regions was calculated as the functional connectivity. The Shen atlas is a functional brain parcellation based on spectral clustering analysis of resting-state fMRI data that has good reproducibility in healthy subjects [13]. We chose this atlas because it has been widely used for CPM analysis and has shown strong predictive performance for various tasks [14-19]. Regions not covered by functional images from all participants due to limited slice coverage or significant signal attenuation, as well as non-gray matter regions, were excluded from the analysis. Supplementary Figure S1 shows the map of the excluded regions. The connectivity between the remaining 233 regions was taken as the whole-brain connectivity pattern. The beta-series FC for the response to the feedback signal was also calculated in the same atlas regions. The interaction between FCs for the regulation task (RegTask) and the response to the NF signal (RespNF) was calculated by multiplying the FC values for each connectivity after standardizing (zero mean, unit variance) each RegTask and RespNF value across all connectivity.

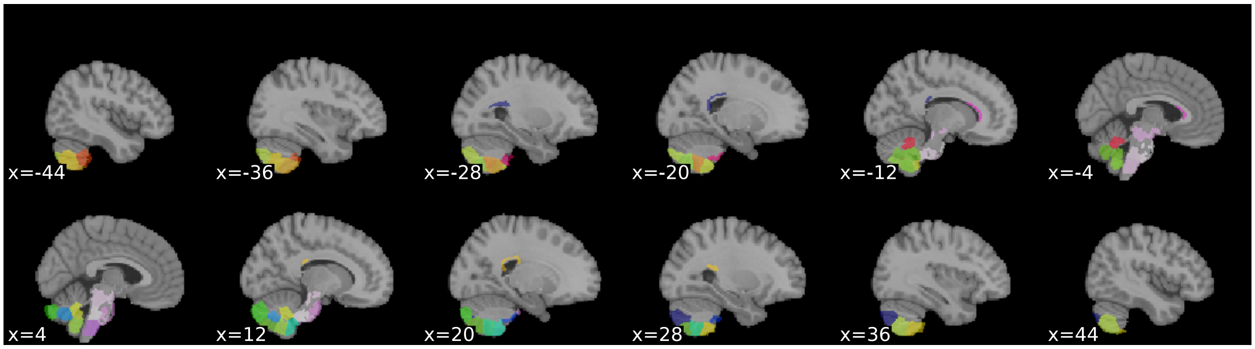

**Figure S1.** Regions excluded from CPM analysis in the Shen 268-node atlas due to limited coverage in the fMRI image. The atlas image was downloaded from https://www.nitrc.org/frs/download.php/7976/shen_1mm_268_parcellation.nii.gz. The indices of the excluded regions were 100, 104, 105, 108, 109, 111, 112, 115, 116, 117, 118, 120, 129, 130, 131, 132, 133, 217, 236, 237, 239, 240, 242, 243, 246, 249, 250, 252, 255, 256, 257, 265, 266, 267, 268.

### Connectome-based Predictive Modeling (CPM)

Connectome-based Predictive Modeling (CPM) is a machine learning approach that builds a predictive model of the brain-behavior relationship from whole-brain functional connectivity patterns [14]. We used CPM to predict the change in symptom score (i.e., RRS-B) one week after the NF session. The prediction target was the ratio of RRS-B change from baseline; dRRS-B = ($\text{RRS-B}_{post}$ - $\text{RRS-B}_{pre}$) / $\text{RRS-B}_{pre}$.

CPM constructed a prediction model using the following steps: 1) selecting connectivity with a high absolute Pearson's correlation ($r$) with the dRRS-B (the $r$ threshold was optimized using nested cross-validation), 2) summing the selected connectivity values for each positively and negatively correlated ones in each participant, and 3) fitting a linear regression model to predict the dRRS-B based on the summed scores of positively and negatively correlated connectivity values, respectively [14]. The prediction model was built for the combined set of active and sham groups. The performance of the prediction model was assessed by Spearman's correlation coefficient between the true and predicted values.

The whole-brain FC patterns of RegTask, RespNF, and their interaction (RegTask:RespNF) were used individually to build the CPM model with multiple combinations. Specifically, we evaluated five CPM models; 1) the main effect of RegTask: dRRS-B ~ RegTask, 2) the main effect of RespNF: dRRS-B ~ RespNF, 3) the sum of these FCs: dRRS-B ~ RegTask + RespNF, 4) the sum of these FCs and their interaction: dRRS-B ~ RegTask + RespNF + RegTask:RespNF, and 5) the main effect of RegTask with the interaction: dRRS-B ~ RegTask + RegTask:RespNF. For example, the CPM model of dRRS-B ~ RegTask + RespNF included four terms (and the intercept) in the linear model, including the sum of positively correlated FCs in RegTask, the sum of negatively correlated FCs in RegTask, the sum of positively correlated FCs in RespNF, and the sum of negatively correlated FCs in RespNF. The model with RegTask:RespNF alone was also tested as a post-hoc evaluation.

Predictive performance was assessed using 5-fold cross-validation, with participants divided into training (80%) and test (20%) groups. The division was made to balance age, sex, head motion, and active/sham members. The covariate effects of age, gender, and head motion (mean FD) were regressed and removed from the individual connectivity and outcome scores (dRRS-B). Covariate regression was performed as part of cross-validation: a covariate regression model was evaluated on the training data, and then the fitted model was used to remove the effects from both the training and test sets. The model hyperparameters, such as the absolute correlation thresholds for connectivity selection, were optimized through a nested 4-fold cross-validation within the training set. The training set was further divided into a training subset (75%) and a validation subset (25%). Grid search was used to explore different values for the hyperparameters: correlation values ($r$) ranging from 0.05 to 0.5 with intervals of 0.05. The entire process of 5-fold cross-validation was repeated 100 times to obtain a confidence interval and a reliable estimate of the predictive performance.

**Table S1**. Top 10 nodes with the largest absolute sum of FC correlation with RRS-B reduction selected in the CPM prediction for FC during the regulation task (RegTask).

| Node | FC correlation | System | Lobe | Gyrus | Hemi | Brainnetome atlas label |
|---|---|---|---|---|---|---|
| 34 | 12.0 | Cingulo-opercular Task Control | Temporal Lobe | STG | R | TE1.0 and TE1.2 |
| 169 | 11.7 | Cingulo-opercular Task Control | Subcortical Nuclei | BG | L | vmPu |
| 20 | 7.9 | Salience | Frontal Lobe | IFG | R | A45c |
| 71 | 7.1 | Visual | Temporal Lobe | ITG | R | A20cv |
| 170 | 6.8 | Subcortical | Subcortical Nuclei | BG | L | dlPu |
| 127 | 5.7 | Subcortical | Subcortical Nuclei | Tha | R | PPtha |
| 253 | 5.6 | Cerebellar | Cerebellum | Cerebellum | L | Left Crus I |
| 264 | 5.1 | Subcortical | Subcortical Nuclei | Tha | L | cTtha |
| 15 | 5.1 | Salience | Frontal Lobe | SFG | R | A8m |
| 11 | 4.9 | Salience | Frontal Lobe | MFG | R | A8vl |

**Node**: region index of the Shen 268-node atlas. **FC correlation**: absolute sum of functional connectivity correlation with RRS-B reduction. The sum was taken for all connectivity selected with significant frequency (> 76%, $p < 0.001$ with Bonferroni correction) in the 100 bootstrap of CPM evaluations. **Hemi**: brain hemisphere.

**Table S2**. Top 10 nodes with the largest absolute sum of FC correlation with RRS-B reduction selected in the CPM prediction for the FC interaction between the regulation task and in response to the neurofeedback signal (RegTask:RespNF).

| Node | FC correlation | System | Lobe | Gyrus | Hemi | Brainnetome atlas label |
|---|---|---|---|---|---|---|
| 30 | 3.5 | Fronto-parietal Task Control | Frontal Lobe | MFG | R | A6vl |
| 11 | 3.2 | Salience | Frontal Lobe | MFG | R | A8vl |
| 126 | 2.5 | Subcortical | Subcortical Nuclei | Hipp | R | cHipp |
| 17 | 2.1 | Default mode / limbic | Frontal Lobe | OrG | R | A12/47l |
| 91 | 2.0 | Salience | Frontal Lobe | PCL | R | A1/2/3ll |
| 156 | 2.0 | Ventral attention | Frontal Lobe | IFG | L | A44op |
| 14 | 2.0 | Fronto-parietal Task Control | Frontal Lobe | PrG | R | A6cvl |
| 170 | 1.9 | Subcortical | Subcortical Nuclei | BG | L | dlPu |
| 253 | 1.9 | Cerebellar | Cerebellum | Cerebellum | L | Left Crus I |

**Node**: region index of the Shen 268-node atlas. **FC correlation**: absolute sum of functional connectivity interaction with RRS-B reduction. The sum was taken for all connectivity selected with significant frequency (> 76%, $p < 0.001$ with Bonferroni correction) in the 100 bootstrap of CPM evaluations. **Hemi**: brain hemisphere.

### Association of other factors with RRS-B changes

The ANOVA table below presents the results of a linear model analysis for RRS-B change (ratio to baseline) with respect to age, sex, head motion (mean FD), self-rating of regulation success (self-rating), duration of positive feedback presentation (PFB-duration), the interaction between self-rating of regulatory success and group, and the interaction between positive feedback duration and group. No significant effects were found for any of these variables.

| | SS | df | *F* | *p* |
|---|---|---|---|---|
| **age** | 0.009 | 1 | 0.172 | 0.682 |
| **sex** | 0.021 | 1 | 0.431 | 0.517 |
| **mean FD** | 0.065 | 1 | 1.313 | 0.262 |
| **self-rating** | 0.129 | 1 | 2.597 | 0.119 |
| **PFB duration** | 0.000 | 1 | 0.002 | 0.961 |
| **self-rating:group** | 0.109 | 1 | 2.193 | 0.151 |
| **PFB duration:group** | 0.003 | 1 | 0.065 | 0.801 |
| **Residuals** | 1.291 | 26 | | |

SS: Sum of Squares, df: degree of freedom, FD: frame-wise displacement, PFB: positive feedback.

### Voxel-wise activation analysis for the association with RRS-B change

Voxel-wise mass univariate analysis was performed on the RegTask and RespNF beta maps and their interaction map to determine whether the voxel-wise activation associated with the treatment effect was present in these responses. We used linear mixed effects (LME) model analysis with the beta maps as the dependent variable and the fixed effects of group (active/sham), run, dRRS-B, age, sex, and participant as a random effect on the intercept. The LME analysis was performed using the lme4 package [20] with the lmerTest package [21] in the R language and statistical computing, and the map of each contrast was computed using the emmeans package. The statistical map of the voxel-wise LME analysis was thresholded at $p < 0.001$ voxel-wise, and then a cluster extent threshold of $p < 0.05$ was applied. The cluster extent threshold was evaluated using AFNI 3dClustSim with an improved spatial autocorrelation function.

LME analyses for the RegTask beta maps showed a significant main effect of dRRS-B in the left lateral occipital region, whose activation was negatively correlated with dRRS-B (Figure S2 in SI). Greater activity in this region during regulation was correlated with greater RRS-B reduction. As this region is associated with visual motion perception, this may suggest that increased attention to the change in the feedback signal was associated with a more effective treatment outcome.

The significant interaction of dRRS-B by run was found in the right superior frontal regions, right precentral gyrus, and right thalamus (Figure S3 in SI). Participants who showed reduced activation across runs in these areas showed a greater reduction in RRS-B. This pattern could suggest that as one perhaps becomes more efficient or familiar with the task, there might be a reduced demand for certain control and cortical activations. It is also plausible that these regional associations could reflect variety of factors, including potential adaptations to the training, rather than solely reflecting changes in emotional state.

Analysis of the beta maps of RespNF and the RegTask:RespNF interaction showed no significant effect on the main effect of dRRS-B and its interactions with Run and Group.

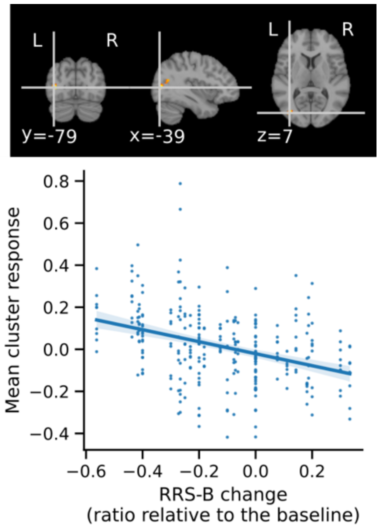

**Figure S2**.

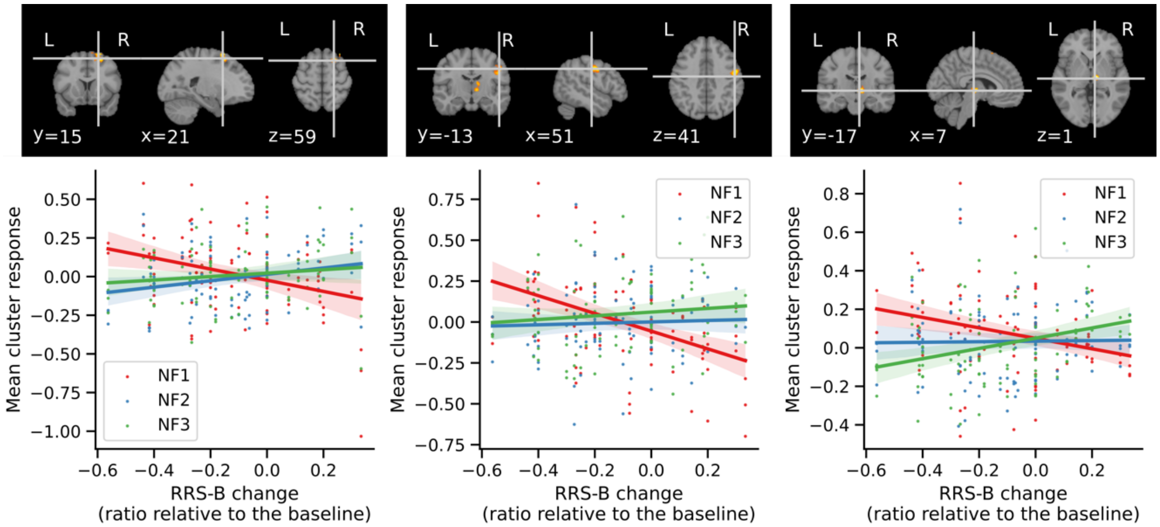

**Figure S3**.
